# Emergence of Pareto Distributions in Intracellular Protein Activity Through Interaction-Driven Modulation

**DOI:** 10.1101/2025.05.16.654514

**Authors:** Naoto Yonekura, Shinji Deguchi

## Abstract

The functional activity of proteins within cells is often unequally distributed: only a small subset of molecules tends to account for the majority of cellular work. This skewed contribution pattern, reminiscent of Pareto’s principle, often known as the 80/20 rule, has been observed across various protein classes, yet its mechanistic origin remains poorly understood. In this study, we present a statistical mechanics-based framework that explains how such disparities naturally emerge from biologically plausible rules of interaction and regulation. By modeling proteins as elements whose activity levels and outputs evolve through mutual comparison and feedback, we demonstrate that power-law distributions can arise without assuming any intrinsic heterogeneity. The model also captures a recursive feature of disparity: even among highly active proteins, a new skewed distribution reappears when a subpopulation is isolated, reflecting the scale-invariant structure commonly observed in complex adaptive systems. We analytically derive these patterns under both positive and negative feedback scenarios and identify key conditions under which long-term functional dominance is established. Our results offer a mechanistic interpretation for the coexistence of active and inactive molecular populations and suggest that functional inequality may reflect an adaptive organizational principle of cellular systems.

## 1. Introduction

Biological systems are inherently hierarchical, with cellular functions emerging from the collective behavior of molecules such as proteins [1]. Protein activity is tightly regulated by conformational changes triggered through interactions with other molecules, including regulatory proteins and substrates. Therefore, protein–protein interactions play a central role in the dynamic regulation of intracellular functions and in the adaptive behavior of living systems.

Interestingly, not all proteins within a cell are active at the same time. Numerous studies have revealed that a substantial proportion of proteins, including functionally critical ones such as ribosomal [2], motor [3], and adhesion proteins [4], remain inactive under physiological conditions. For example, ribosomal proteins, which mediate protein synthesis, are known to be partially inactive in both proliferating and non-proliferating cells, with the inactive fraction increasing significantly in quiescent states [2]. Similarly, only a limited number of myosin [3] and integrin [4] molecules sustain intracellular tension at any given time, despite the presence of a large reserve pool [5]. These findings suggest that cells maintain a functionally redundant set of proteins in order to respond adaptively to environmental changes arising from diverse intracellular and extracellular stresses [6–9].

The selective activation of only a subset of available proteins implies that molecular activity is not distributed uniformly, but is instead shaped by structured, non-random mechanisms such as phosphorylation, complex formation, ubiquitination, acetylation or methylation, and subcellular localization. These regulatory processes give rise to multilayered levels of activation, where proteins are not simply switched on or off, but instead modulate their functional engagement in graded and context-dependent ways such as changing their location within the cell or altering interaction specificity [2–5]. This consistent coexistence of active and inactive molecular populations may reflect a form of functional heterogeneity that is not incidental. Given that proteins can exhibit a wide range of activation states, it is plausible that their functional contributions are not distributed evenly but rather follow a systematic pattern. This raises the possibility that molecular roles within cells are described by an underlying statistical principle that shapes the overall distribution of activity levels.

A well-known example of such a principle is Pareto’s law, originally formulated in economics to describe situations where a small fraction of elements accounts for a large share of the total outcome—a pattern often referred to as the 80/20 rule [10,11]. In living cells, for example, around 30 percent of ribosomal proteins remain inactive even during active cell growth, rising to over 50 percent in quiescent states [2]. While these ratios do not exactly match the classic 80/20 split, they indicate a clear deviation from uniformity, supporting the idea that only a limited subset of molecular components contributes significantly to cellular function. Similar skewed distributions have been documented in diverse complex systems, including ecological networks, computer systems, and social organization [10,11], suggesting that such patterns may arise from shared underlying principles. These observations support the view that such distributions are not mere empirical anomalies, but rather statistical regularities shaped by system-level constraints and interaction dynamics.

Statistical mechanics provides a useful framework for analyzing how macroscopic statistical patterns emerge from microscopic interactions [12]. Rawlings et al. [13] introduced an insightful statistical mechanics-based model, in which Pareto’s law emerges from a quasi-equilibrium ensemble where the sum of the logarithms of individual incomes is conserved. This yields a canonical distribution over log-transformed variables and naturally leads to a power-law distribution. However, this formulation relies on assumptions that are not consistent with the biological reality of molecular systems. Specifically, the elements are treated as non-interacting, and their functional capacities are assumed to remain fixed over time. In contrast, within the molecular complexity of living systems, the capacity of a molecule to perform its function typically varies over time as a result of interactions with other components.

In this study, we describe a theoretical framework that demonstrates how Pareto-like distributions can emerge even when biologically realistic features, such as activation and inactivation governed by molecular interactions, are taken into account. By incorporating interaction-dependent mechanisms of regulation, our model shows that power-law behavior can arise naturally under plausible intracellular conditions. Moreover, the framework captures a form of scale-invariant disparity. Specifically, even when the most active subset of molecular components is extracted, a similarly skewed distribution re-emerges. This invariance under subsystem extraction suggests a fundamental organizational property of cellular systems that may underlie their capacity for adaptive behavior.

## 2. Model

Our model is based on the idea that molecular components do not act in isolation but continuously adjust their functional capacity in response to their interaction and relative state within the population. Such interaction-driven modulation is consistent with known biological mechanisms, including conformational changes and signaling feedback. By capturing the essential regulatory logic of these interdependencies, our approach offers a biologically grounded explanation for the emergence of Pareto-like distributions in molecular systems. Detailed derivations of the equations in this section can be found in Supplementary Information.

### 2.1 Statistical mechanics model of effective molecular contributions

We consider a closed system composed of *N* elements, assumed to be in thermodynamic equilibrium. Each element *i* is characterized by a quantity *u*_*i*_, which represents its effective functional contribution to the system. In biological terms, *u*_*i*_ represents the emergent level of functional engagement resulting from the cumulative effects of multiple regulatory mechanisms such as conformational changes, subcellular localization, and post-translational modifications. In the steady state, the total effective contribution *U* from all elements is assumed to be conserved:

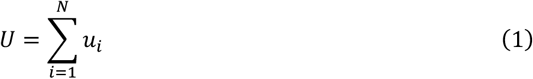

We assume that each element possesses *M* independent features that determine its microscopic state; here, each “feature” may correspond to biologically relevant attributes such as post-translational modifications, conformational states, subcellular localization, or molecular binding partners, all of which influence how each element engages in its biological function. Let 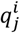 denote the *j*-th feature of element *i*, and let the full state vector of element *i* be defined as

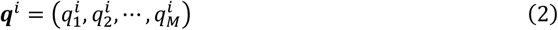

The quantity *u*_*i*_ is then assumed to be a function of this feature vector:

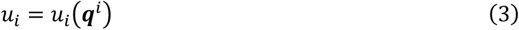

We invoke the principle of equal a priori probabilities, which states that in a closed system at equilibrium, all microstates consistent with the macroscopic constraints are equally probable. Specifically, for any two elements *i* and *j*, if their state vectors are identical (i.e., ***q***^*i*^ = ***q***^*j*^), then the probability that element *i* occupies state ***q****i* is equal to that for element *j*. Under this assumption and given the freedom for elements to exchange *u*_*i*_ within the closed system, the probability *p*_*i*_ that an element occupies state ***q***^*i*^, namely possesses *u*_*i*_(***q***^*i*^), follows the canonical distribution:

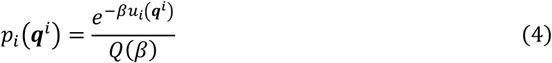

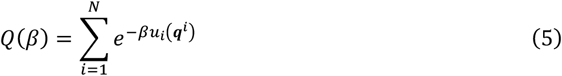

Here, *β* represents the inverse temperature of the system, a macroscopic parameter characterizing the equilibrium state.

### 2.2 Time-dependent modulation of protein output and activity

We consider a system composed of *N* elements representing proteins of interest within a cell. Each protein *i* is characterized at time *t* by two state variables: its functional output, denoted by *z*_*i*_(*t*), and its activity level, denoted by *ε*_*i*_(*t*). The functional output *z*_*i*_(*t*) represents the realized workload of the protein, such as driving the synthesis of other proteins, generating mechanical forces, or maintaining cellular architecture. The activity level *ε*_*i*_(*t*), on the other hand, describes the functional capability of the protein, which may vary over time due to factors such as conformational changes, post-translational modifications, or changes in subcellular localization. Unlike the realized functional output *z*_*i*_(*t*), which reflects the actual work performed, or the effective functional contribution *u*_*i*_, which quantifies the system-level impact of each protein, *ε*_*i*_(*t*) reflects the functional potential of the protein to carry out its role, which can be dynamically modulated by intermolecular interactions. We assume that proteins are engaged in some level of activity; hence, the functional output *z*_*i*_(*t*) is bounded below by a positive constant *δ* (> 0).

The time evolution of *z*_*i*_(*t*) is modeled by a differential equation that reflects two key factors: the current output level and the relative difference between the present activity of the protein of interest and the average activity of the other proteins in the system. Specifically, we define

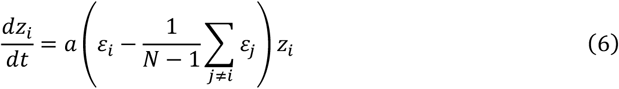

where *a* is a proportionality constant. This formulation captures the intuitive idea that the output of a protein increases or decreases depending on whether its activity level is above or below the population average. It also reflects the notion of competition or interaction among proteins.

In contrast, the time evolution of the activity level *ε*_*i*_(*t*) is left unspecified and may be governed by an arbitrary function. This is a deliberate choice, as protein activity level depends on complex biochemical and mechanobiological pathways and is shaped by the conformational states of the protein. In some biological contexts, activity level may increase as a result of accumulated output, such as when sustained functional engagement leads to cooperative interactions that enhance molecular performance. In other contexts, continued output may instead lead to a decline in activity level, for example, due to molecular fatigue, resource depletion, or inhibitory post-translational modifications, resulting in the downregulation of functional engagement despite ongoing contribution.

### 2.3 Output distributions emerging from protein interactions

Using Eqs. (4) and (5), we derive the probability that a protein exhibits a given output *z*_*i*_. To do so, the functional form of the effective contribution *u*_*i*_ (*z*_*i*_, *ε*_*i*_) must first be determined. The total activation across all proteins is given by *U*, which remains constant over time at the steady state. Thus, its time derivative satisfies

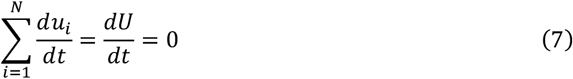

Meanwhile, from the time evolution of the output in Eq. (6), we can derive the following identity involving the logarithmic derivative of *z*_*i*_(*t*):

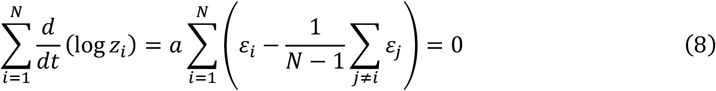

Comparing Eqs. (7) and (8), we deduce that the effective contribution must take the form

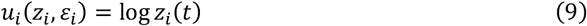

Accordingly, the joint probability that a protein exhibits a given level of output *z*_*i*_ and activity level *ε*_*i*_ can be expressed using Eqs. (4) and (5):

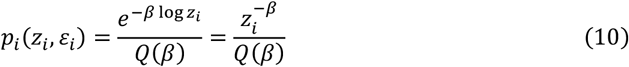

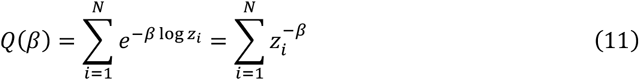

By treating *z* and *ε* as continuous variables, we derive the joint probability density function *f*(*z, ε*) from Eqs. (10) and (11). Here, the domains of *z* and *ε* are taken to be the same as those of *z*_*i*_ and *ε*_*i*_, respectively.

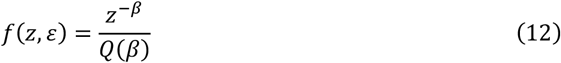

where the normalization factor is given by

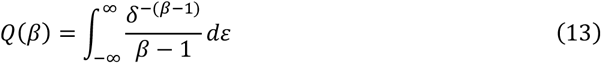

This result holds under the condition *β* > 1. The marginal distribution for the output *z* is then given by

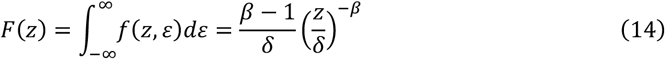

This is a Pareto distribution, confirming that the functional output distribution derived from our model conforms to the Pareto law. Furthermore, the cumulative probability *G*(*z*) that the output exceeds a given threshold *z* takes the power-law form

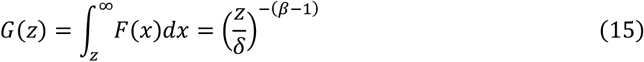

In order for the probability density function *F*(*z*) to follow a Pareto distribution of the form *F*(*z*) *∝ z*^−(*γ* +1)^ for any *γ* > 0, and given that *F*(*z*) *∝ e*^− *βu*(*z*,*ε*)^, it must follow that

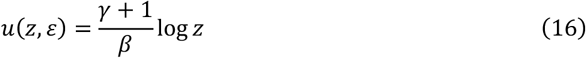

This matches the functional form in Eq. (9), which results directly from the interaction-based constraint in Eq. (8). In particular, Eq. (8) captures how the capacity of each protein is compared to the population average, reflecting a fundamental interaction structure within the system. Such dynamics are biologically plausible, as proteins continuously regulate one another through molecular interactions. Our model thus reproduces the Pareto distribution under biologically meaningful assumptions and highlights the essential role of protein interactions in shaping system-level statistical patterns.

### 2.4 Feedback-driven enhancement of protein activity

In many biological contexts, the conformational state and functional efficiency of a protein are regulated through interactions with other molecular species. Such regulatory mechanisms are generally referred to as allosteric control, in which the binding of a protein to a distinct effector molecule induces a structural transformation, altering its functional activity. For example, in a biochemical pathway, the initial enzyme may bind to the final product of the same pathway, resulting in either an enhancement or suppression of its activity. Thus, allosteric regulation can be either positive or negative. In this section, we focus on the former scenario, wherein the activity of a protein is upregulated as a consequence of its own prior activation, a situation commonly observed in feedback-driven systems within cells.

We assume that the activity level *ε*_*i*_ of protein *i* evolves over time in response to its own past activation relative to that of other proteins in the system. This dependence is modeled as

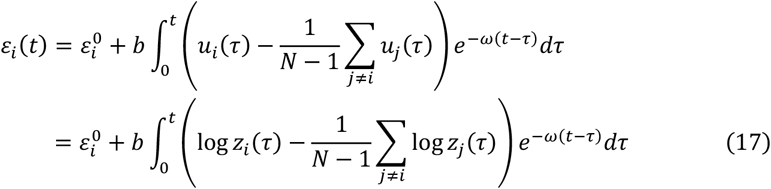

Here, 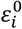 denotes the initial value of *ε*_*i*_(*t*), while *b* and *ω* are positive constants. The parameter *b* characterizes the strength of influence that *u*_*i*_(*t*) has on *ε*_*i*_(*t*), and *ω* controls the rate at which the influence of past contribution decays over time. The integral on the right-hand side represents a convolution between the temporal deviation of *u*_*i*_(*t*) from the population mean and an exponential decay kernel *e*^−*ω* (*t*−*τ*)^, capturing the biologically plausible assumption that more recent contribution has greater impact on present activation.

By analytically solving Eq. (6), which describes the time evolution of the functional output *z*_*i*_(*t*), in conjunction with Eq. (17), we obtain an explicit expression for the trajectory of *z*_*i*_(*t*) under a feedback-driven regulation scheme as follows:

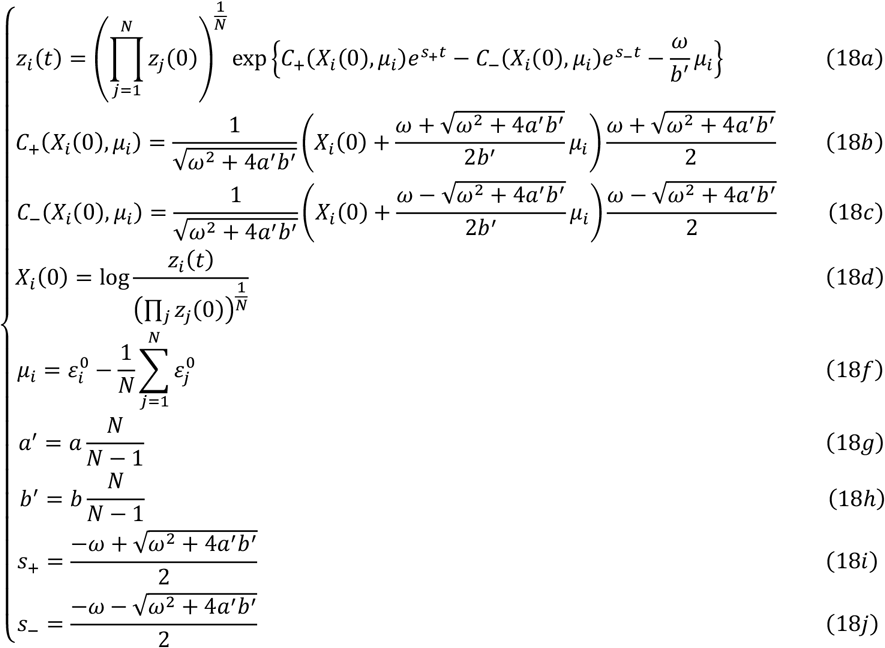

In Eq. (18a), the parameters *s*_+_ and *s*_−_ represent the exponential growth and decay rates, respectively, of the output *z*_*i*_(*t*). By construction, *s*_+_ is always positive and *s*_−_ is negative. Therefore, in the long-time limit, the first term within the braces of Eq. (18a) dominates the behavior of *z*_*i*_ (*t*), and the output can be approximated as

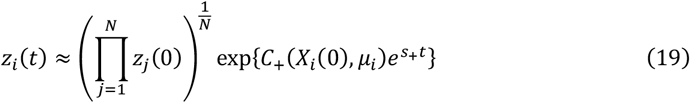

This expression implies that, as time progresses, the long-term behavior of *z*_*i*_ (*t*) is governed by the sign of the coefficient *C* _+_ (*X*_*i*_(0), *μ*_*i*_). If *C*_+_ > 0, the functional output of protein *i* increases exponentially and becomes large; conversely, if *C*_+_ < 0, the output diminishes and approaches a negligible value. Thus, the sign of *C*_+_ has a decisive impact on the eventual activity level of the protein. By introducing the quantity

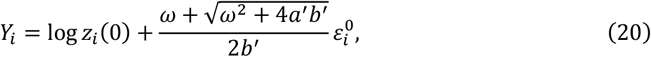

we find that the condition under which protein *i* becomes highly active in the long-time limit is

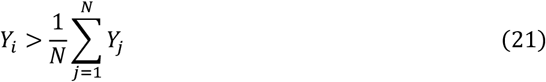

This result indicates that the relative value of *Y*_*i*_, determined by the initial output *z*_*i*_(0) and the initial activity level 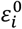, predicts whether a protein will become highly active within the population. In other words, *Y*_*i*_ serves as a key quantity that governs the long-term functional role of protein *i* in the system. Furthermore, if we isolate the subset of proteins that are highly active and form a new subpopulation, the average value of *Y*_*i*_ within this subpopulation will exceed that of the original population. Because the condition for high activity depends on comparison with the population mean, this recursive selection naturally leads to the emergence of less active proteins within any new group (Fig. 1).

**Fig. 1.**
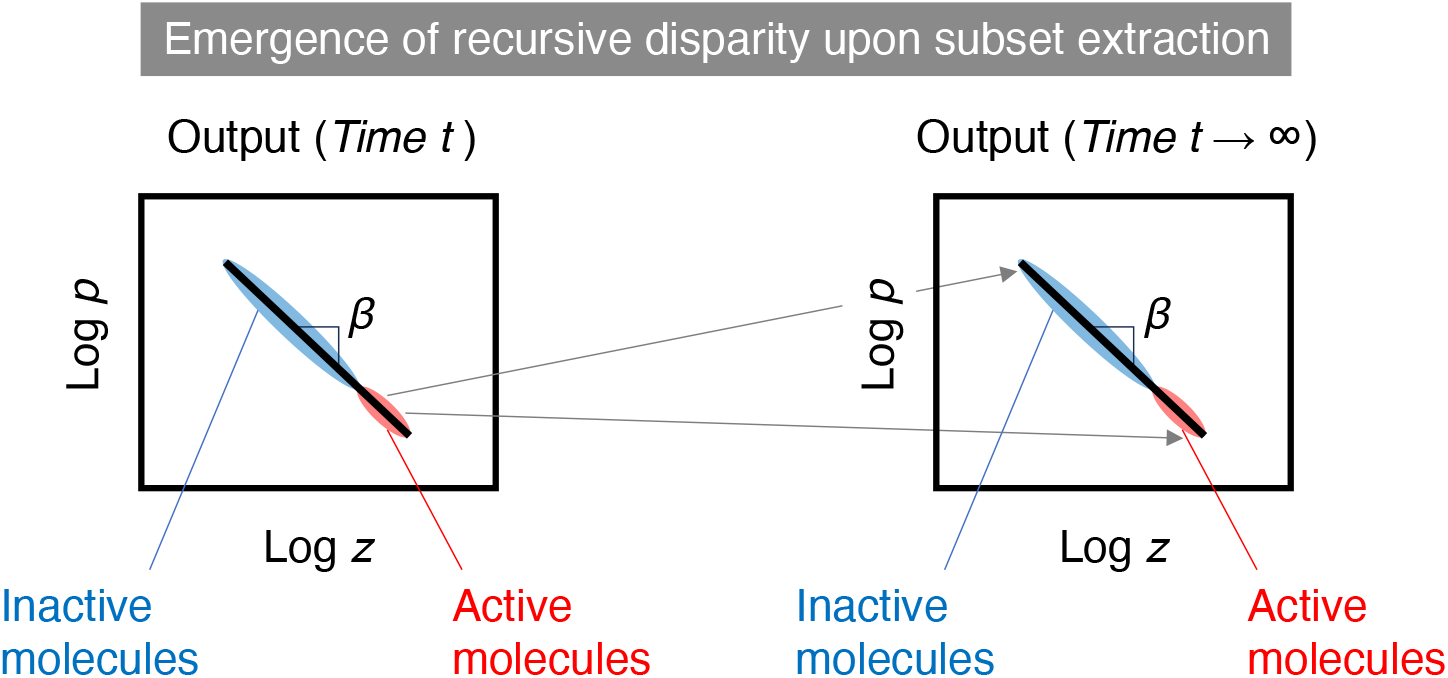
Pareto-like behavior emerging from the present model.

### 2.4 Feedback-driven suppression of protein activity

Allosteric regulation can involve either positive or negative control, depending on whether molecular binding enhances or suppresses protein function. While the previous section focused on a model of positive regulation, where activity promotes further activation, we now turn to the case of negative regulation, in which the functional capacity of a protein is diminished as a consequence of its own activity. Following the same modeling framework as before, we describe the time-dependent activity level *ε*_*i*_ of protein *i* under negative feedback as

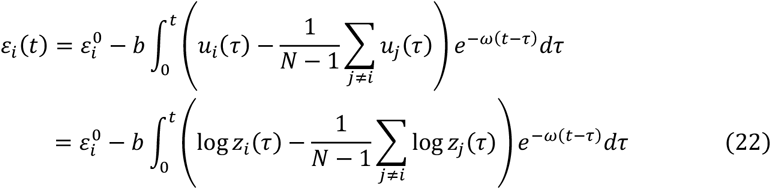

This equation differs from the positive feedback case in Eq. (17) only in the sign of the second term. The negative sign indicates that increased activation leads to a decline in the activation level of the protein, reflecting inhibitory regulation.

As in the previous section, we analytically solve the coupled differential equations composed of the output dynamics in Eq. (6) and the inhibitory activity dynamics in Eq. (22) using Laplace transform techniques. Explicit solutions for the time evolution of the workload of each protein under negative feedback were then obtained:

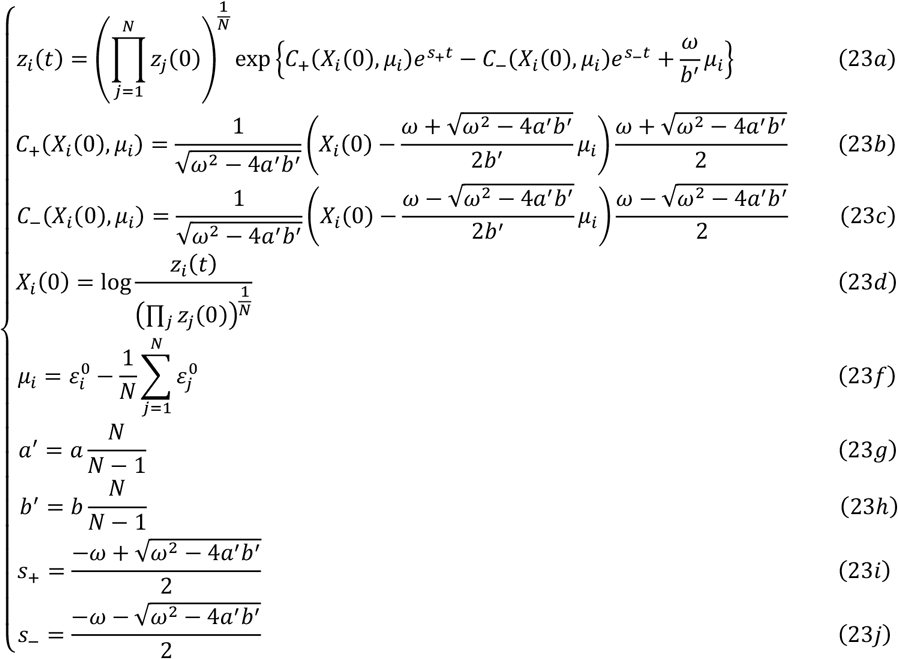

The long-term behavior of *z*_*i*_(*t*) is shaped by the sign of the discriminant *ω*^2^ − 4*a*^′^*b*^′^, which determines whether the system exhibits monotonic or oscillatory dynamics. Because *ω*, *a*^′^, and *b*^′^ are all positive real parameters, two qualitatively distinct regimes arise depending on whether *ω*^2^ is greater or less than 4*a*^′^*b*^′^. When *ω*^2^ > 4*a*^′^*b*^′^, the roots *s*_+_ and *s*_−_ that govern the dynamics of the system are both real and negative. In this regime, the exponential terms in Eq. (23a) decay over time, rendering their influence negligible in the long-time limit. As a result, *z*_*i*_(*t*) approaches a steady-state value that is determined solely by the initial conditions

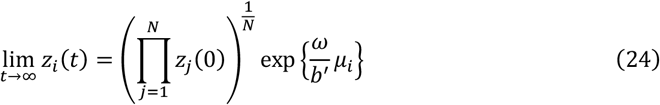

This expression indicates that the steady-state level of functional output is determined by *μ*_*i*_. Proteins with higher *μ*_*i*_ values tend to exhibit higher long-term contribution, highlighting the importance of early molecular states in shaping later functional outcomes.

In contrast, when *ω*^2^ < 4*a*^′^*b*^′^, the roots become complex conjugates and the functional output exhibits transient oscillations. In this case, the dynamics of *z*_*i*_ (*t*) are governed by a combination of decaying exponential and sinusoidal components

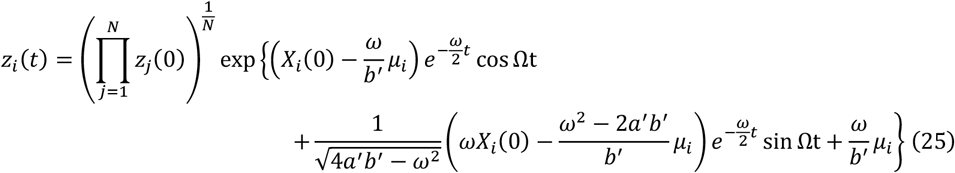

where the oscillation frequency Ω is given by

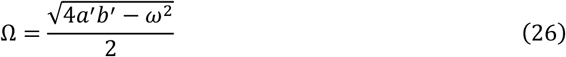

Although the trajectory of *z*_*i*_(*t*) in this regime is oscillatory, the amplitude of the oscillations diminishes with time, and the output eventually converges to the same steady-state value as in Eq. (24). Thus, regardless of whether the dynamics are monotonic (*ω*^2^ > 4*a*^′^*b*^′^) or oscillatory (*ω*^2^ < 4*a*^′^*b*^′^), the long-term behavior of each protein depends on its initial capacity relative to the population. This leads to a similar conclusion as in the positive regulation model in that proteins with relatively high initial activity become highly active in the long term. Moreover, when only such highly active proteins are selected to form a new population, the population average of initial activity increases, and new disparities emerge (Fig. 1). Some proteins in the new group will again fall below the new mean and appear less active by comparison. Thus, even in this inhibitory feedback model, the system exhibits a recursive structure, in which functional disparity persists regardless of how the population is selected or redefined.

## Discussion

Understanding how unequal molecular activity emerges within cells is essential to deciphering the statistical architecture of intracellular function. In this study, we introduced a statistical mechanics-based model that accounts for time-dependent molecular activity, interaction-driven regulation, and system-level feedback.

Unlike earlier approaches [13], which assumed fixed capabilities and independence among elements, our formulation reflects the biological reality that proteins continuously modulate their activity in response to environmental cues and interactions with other molecules. This model introduces a distinction among three variables: functional output *z*_*i*_(*t*), activity level *ε*_*i*_(*t*), and effective functional contribution *u*_*i*_(*z*_*i*_, *ε*_*i*_). *ε*_*i*_(*t*) represents the evolving capability of a protein to perform its function, influenced by regulatory mechanisms such as conformational changes, localization shifts, and post-translational modifications. *z*_*i*_(*t*) refers to the realized functional expression, meaning the actual work performed by the protein, while *u*_*i*_ reflects the cumulative contribution of the protein to the system.

We analytically derived a power-law distribution of functional output from an interaction-driven framework, consistent with observations in various biological systems [2-4,14,15]. The key lies in a comparison mechanism: the change in output depends not only on absolute activity but on its relative standing within the population. This comparative structure introduces a hierarchy among otherwise identical components, naturally producing disparities in output. Because functional impact evolves over time through feedback and relative evaluation of activity, proteins with initially similar capacities can diverge significantly. Importantly, the model predicts that such disparities recur even when only the most active proteins are selected, giving rise to recursive disparity, a scale-invariant structure characteristic of complex systems [10, 16, 17]. These findings suggest that functional inequality is not an incidental outcome but a structurally embedded feature of cellular organization.

This theoretical prediction aligns with empirical evidence: many proteins, such as ribosomal, motor, and adhesion proteins [2–5], are frequently found in inactive states under physiological conditions, despite their essential roles. Our results provide a mechanistic interpretation of this observation, suggesting that the presence of low-output proteins may not reflect inefficiency but instead contribute to cellular robustness. These proteins can serve as a latent reserve, allowing cells to rapidly adapt under stress or perturbation when increased demand for function arises [6-9]. Importantly, our model also exhibits recursive disparity, a structure analogous to recurring inequalities that have been noted in various complex systems such as economic, ecological, social networks [16,17]: i.e., even when only the most active proteins are selected, a new skewed distribution re-emerges within the subset. This scale-invariant behavior reflects a fundamental organizational feature of intracellular systems, suggesting that functional inequality is not incidental but structurally embedded through feedback and comparative regulation.

While our model successfully captures the emergence of power-law distributions in the high-output regime, it does not account for the distributional behavior observed at lower output levels. Experimental studies have reported that protein activity can follow a log-normal distribution in this range, with a crossover to power-law behavior only above certain thresholds [13]. Capturing this dual behavior remains an important direction for future theoretical refinement. Furthermore, the present model is built on mean-field comparisons, assuming global evaluation of relative activity. Incorporating the actual topological features of molecular interaction networks, including local interactions, modularity, and heterogeneity, may reveal additional layers of disparity and provide a more nuanced understanding of intracellular regulation.

In conclusion, this study offers a mechanistic perspective on the unequal distribution of protein activity observed in cells. By incorporating regulatory comparisons and feedback into a statistical mechanics framework, we show that power-law distributions and recursive disparities can arise naturally from local molecular interactions. These results contribute to a broader understanding of how cells organize their functional architecture, not through centralized coordination, but through simple statistical rules embedded in distributed interaction networks.

## Acknowledgments

This study was partly supported by JSPS KAKENHI grants (21H03796).

## Declaration of interests

The authors declare no competing interests.

## Supporting Information

Here, we provide a supplementary explanation regarding the derivation of the equations presented in the main text. Regarding Eq. (8), given that the functional output *z*_*i*_ (*t*) remains strictly positive and that its time evolution is governed by Eq. (6), the following identity for *z*_*i*_ (*t*) holds:

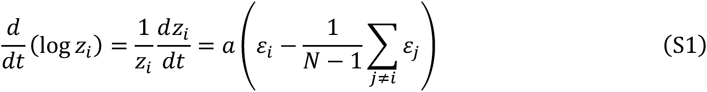

By summing this equation over all indices *i*, we obtain Eq. (8).

A more detailed derivation of Eq. (13) is presented as follows:

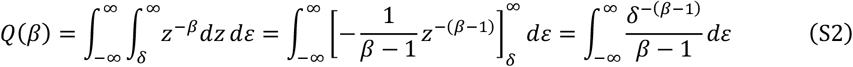

From this expression, it follows that *β* > 1 is required to ensure the convergence of the integral.

The detailed derivations of Eqs. (14) and (15) are provided below:

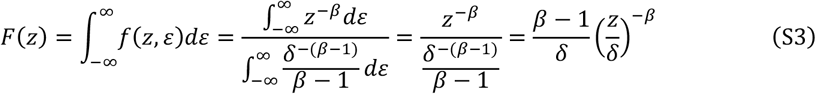

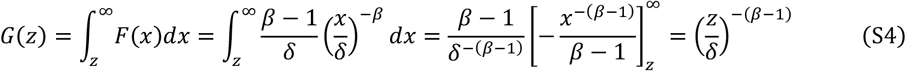

We provide additional details on the analytical solution of the coupled differential equations governing the time evolution of *z*_*i*_ (*t*) and *ε*_*i*_ (*t*) described by Eqs. (6) and (17), respectively. We begin with

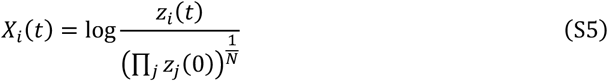

which, together with Esq. (S1) and (S5), allows us to obtain the following:

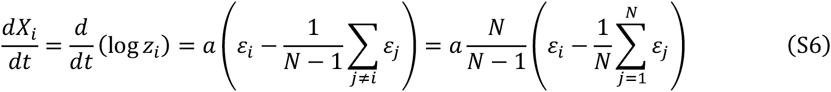

Then, from Eq. (17),

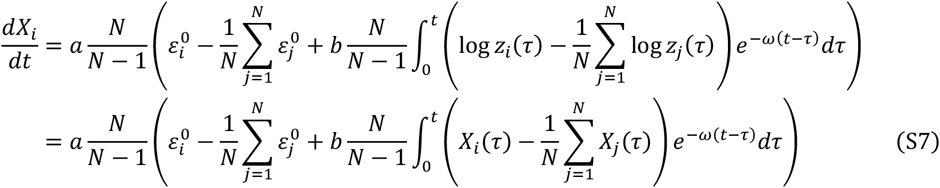

By summing Eq. (S7) over all indices *i*, we obtain the following:

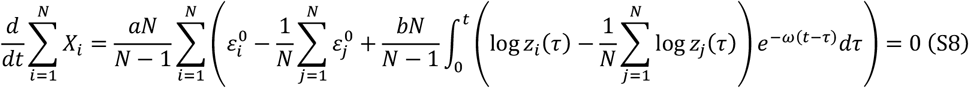

The total sum of *X*_*i*_ (*t*) is then given by

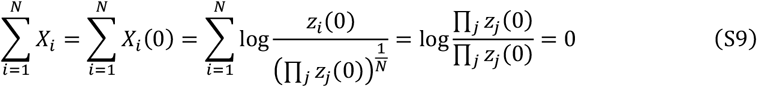

Therefore, Eq. (S7) is expressed as

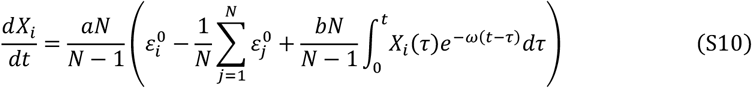

We define the Laplace transform of *X*_*i*_ (*t*), denoted 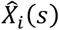 as follows:

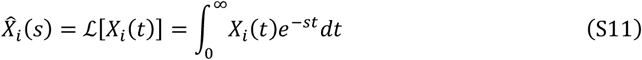

where *s* is a complex variable. For notational convenience, we define the following expressions:

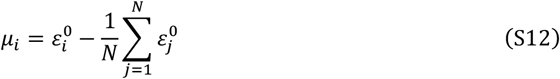

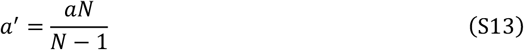

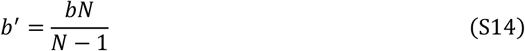

Applying the Laplace transform to Eq. (S10) yields the following:

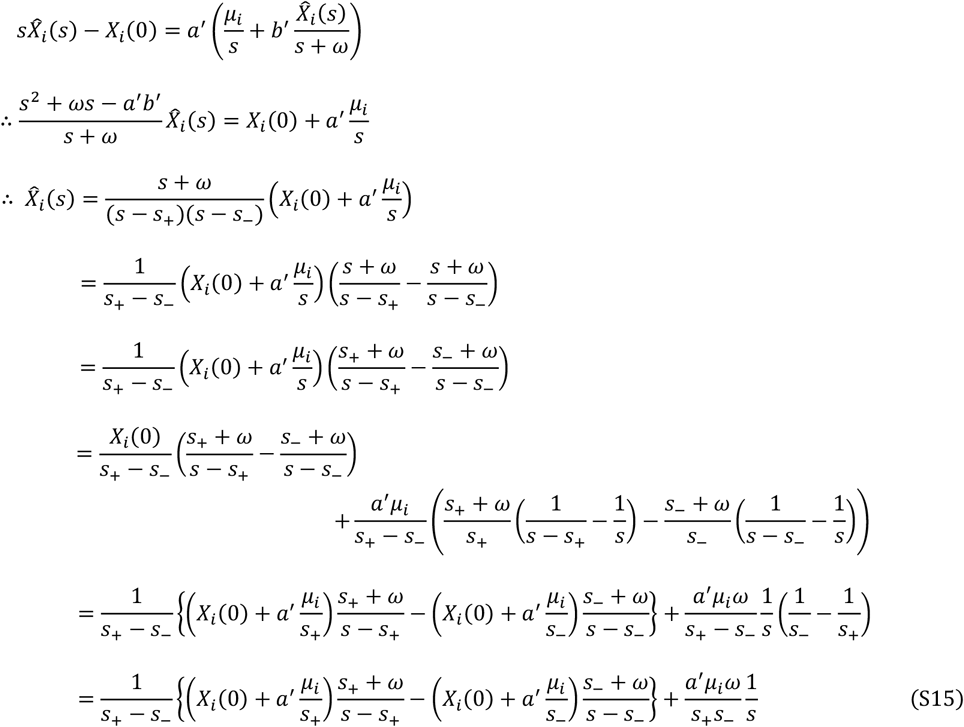

where *s*_+_ and *s*_−_ are the solutions of the following equation

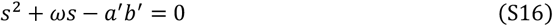

and thus,

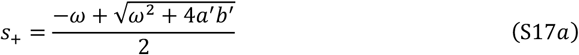

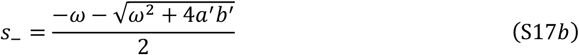

Hence,

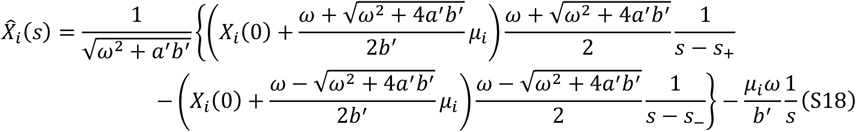

By applying the inverse Laplace transform, we analytically obtain the functional output or workload of individual proteins *z*_*i*_(*t*) as in Eq. (18).

We now consider the condition under which *z*_*i*_(*t*) of a protein becomes large in the long-time limit. In the limit *t* → ∞, for *z*_*i*_(*t*) → ∞, the coefficient *C*_+_(*X*_*i*_(0), *μ*_*i*_) must be positive. From Eq. (18b), this condition can be derived as follows:

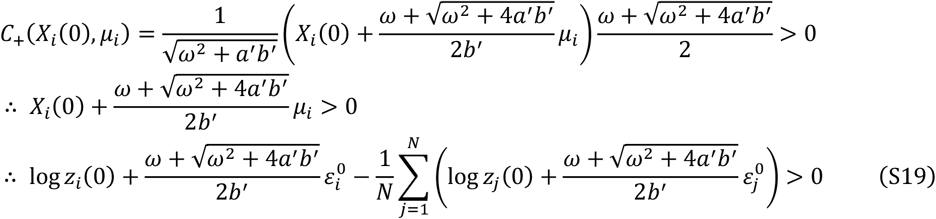

By introducing the quantity *Y*_*i*_ defined in Eq. (20), the condition in Eq. (21) can be equivalently expressed. The time evolution of *z*_*i*_(*t*) is illustrated in Fig. S1.

**Fig. S1.**
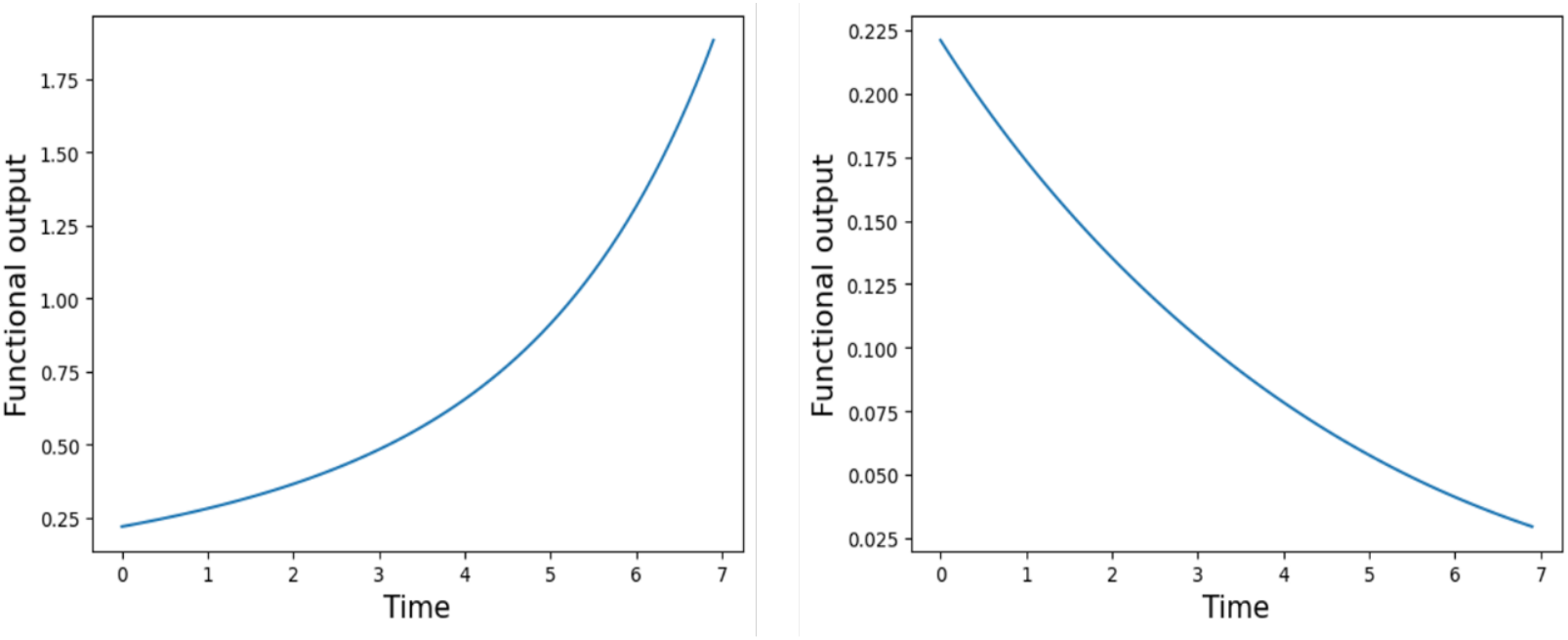
Workload (functional output) of *i*th protein. (a) A representative case in which the protein becomes highly active in the long-time limit. *N* = 100, *a* = 0.24, 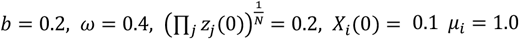. (b) A case in which the protein becomes inactivated over time. *N* = 100, *a* = 0.24, *b* = 0.2, 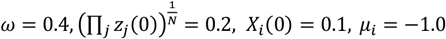.

Next, we derive the time evolution of *z*_*i*_(*t*) in the case where *ω*^2^ < 4*a*^′^*b*^′^. In this regime, the coefficient *C*_+_(*X*_*i*_(0), *μ*_*i*_) takes the following form, in which the subscript *i* denotes the index of the protein, whereas any other occurrence of *i* refers to the imaginary unit.

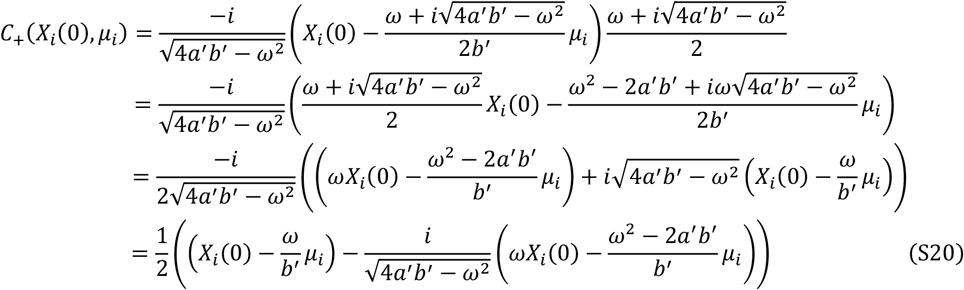

Likewise,

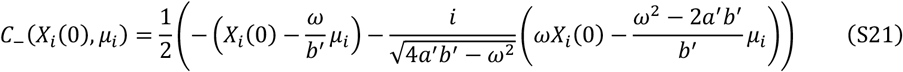

Let us define the following quantities:

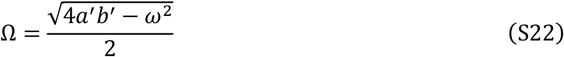

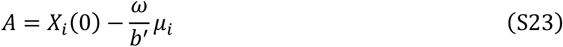

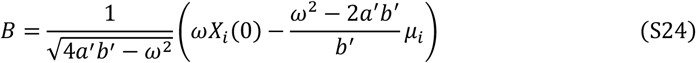

The first term inside the braces in Eq. (23a) is described by

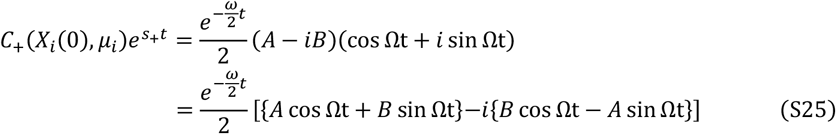

In the same manner, the second term inside the braces is

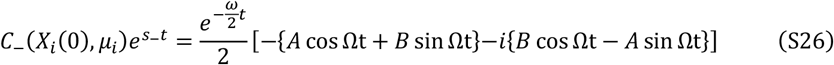

Thus, the functional output of the protein takes the following form:

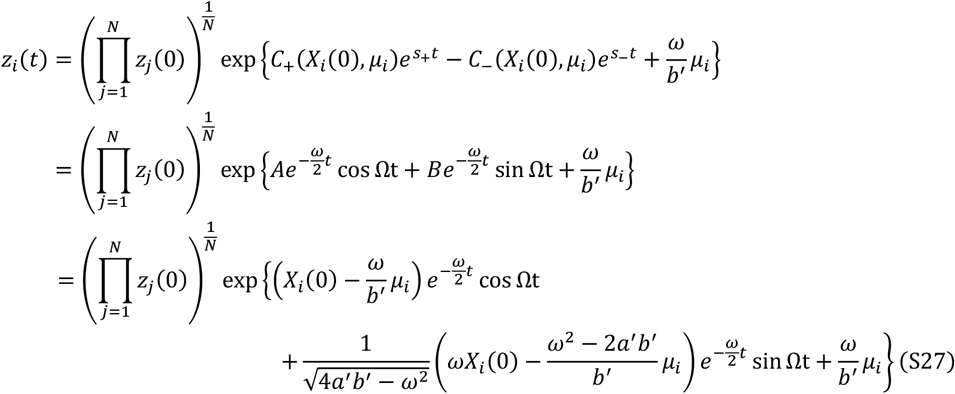

The time evolution of *z*_*i*_(*t*) for individual proteins is illustrated in Fig. S2.

**Fig. S2.**
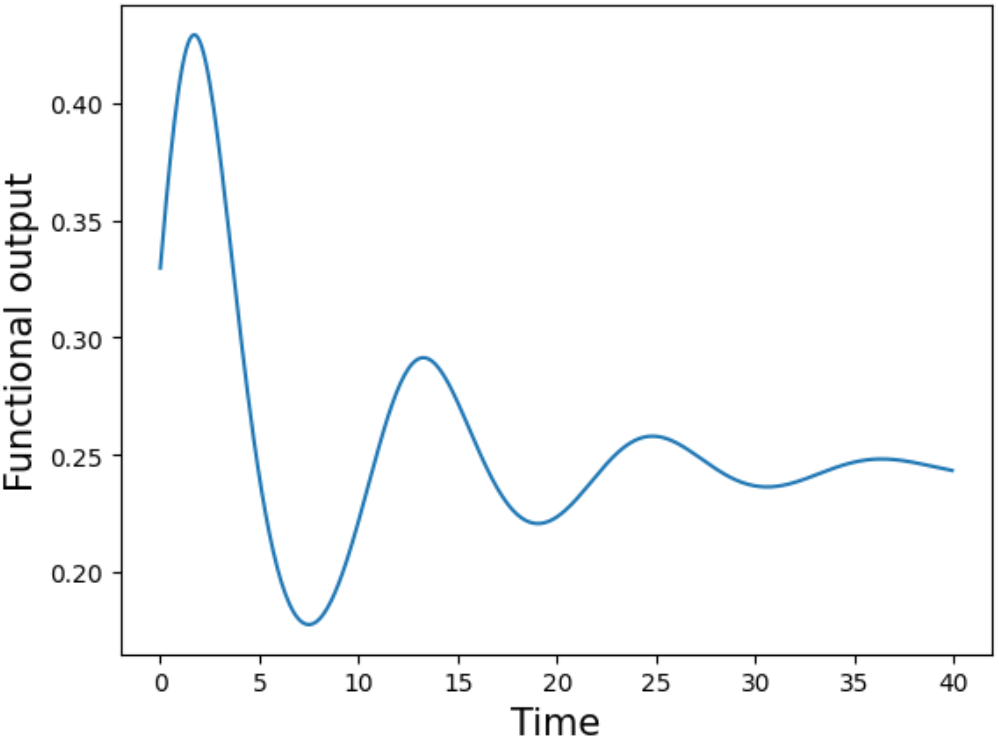
Workload (functional output) of *i* th protein. *N* = 100, *a* = 0.6, *b* = 0.5, 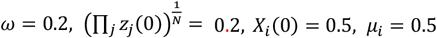.

## Notes

### Competing Interest Statement

The authors have declared no competing interest.

